## Supplementary material for "Controlling gene expression timing through gene regulatory architecture": S1 Text

### S1 Text: Cell-cycle effect

In the manuscript, we have assumed that cells dilute protein through random degradation at a rate $ln(2)/\tau$, where $\tau$ is the cell division time. However, the proteins can be very stable and the way it is diluted is mainly through cell division where the cell contents are roughly equally allocated between the mother and the daughter cells. In order to show that even when the proteins are diluted purely through cell division all the results discussed in the manuscript is preserved, we model the process of cell division exclusively in our stochastic simulations using the same reaction schemes described in the Materials and Methods section. The cell division time is chosen from a Gaussian distribution with mean time ($\tau_{\mathrm{c}}$) and a standard deviation ($\sigma_\tau$), every time the cell divides. When the cell divides, proteins and mRNAs are partitioned binomially between the mother and daughter cells and the mother cell is kept. For simplicity, we do not model gene replication and keep the gene copy number one all the time. If the gene is occupied by a TF then after cell-division it remains occupied with a probability set by the total binding rate ($n_{\mathrm{TF}}k_{\mathrm{on}}$) and the off-rate ($k_{\mathrm{off}}$). If the gene is not bound then it remains unbound even after the cell divides. In Fig 1, we show that all the results are conserved for auto-activating and auto-repressing TF gene when off-rate is tuned.

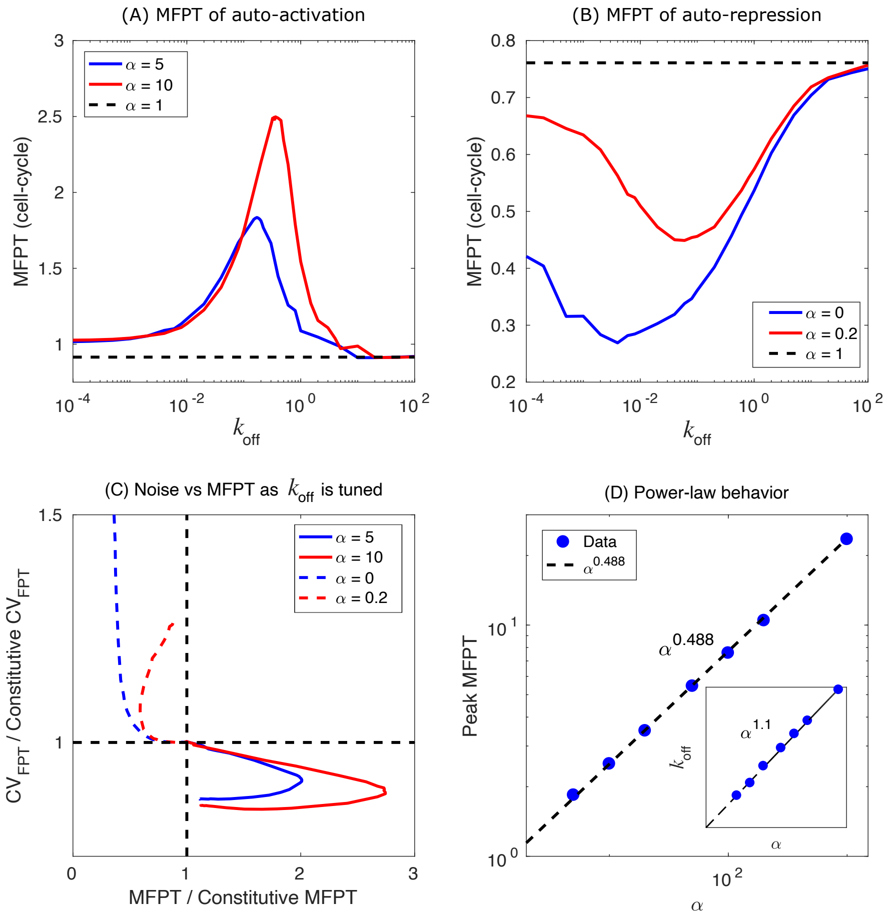


**Fig 1. Simulating cell-cycle in the stochastic model.** MFPT of an auto-activated (A) and and auto-repressed (B) TF gene as a function of TF off-rate ($k_{\text{off}}$). The dashed lines correspond to MFPT of a constitutive gene. (C) The CV of first passage time as a function of MFPT when koff is changed to vary MFPT for auto-activation (solid lines) and auto-repression (dashed lines) of differing regulatory strength. (D) Peak value of MFPT as a function of auto-regulation strength, $\alpha$, when $k_{\text{off}}$ is tuned (circles). The dashed line is a power-law fit with an exponent 0.488. (Inset) The values of $k_{\text{off}}$ at the peak MFPT as a function of $\alpha$ also follow a power-law with an exponent 1.1. Parameters: $\gamma_{\text{m}} = 0.01\ \text{s}^{-1}\text{mRNA}^{-1}$, $\alpha = 5,\ 10$, $r_0 = 0.0025\ \text{s}^{-1}$, $b = 0.025\ \text{s}^{-1}$ (auto-activation), $\alpha = 0,\ 0.2$, $r_0 = 0.01\ \text{s}^{-1}$, $b = 0.1\ \text{s}^{-1}$ (auto-repression). Cell-division time is drawn from a Gaussian distribution (mean, $\tau_{\text{c}} = 50$ min and standard deviation, $\sigma_\tau = 5$ min. In addition to dilution through cell-division, protein degrades with a half life of 5 hours.
