## Supplementary material for "Controlling gene expression timing through gene regulatory architecture": S2 Text

#### S2 Text: Deterministic Approach

The reactions described in Materials and methods section can be translated into a set of ordinary differential equations (ODEs) using mass action kinetics and is given by

$$\begin{aligned}
\frac{dn_{\mathrm{TF}}}{dt} &= bm_{\mathrm{TF}} - \gamma n_{\mathrm{TF}} - k_{\mathrm{on}} n_{\mathrm{TF}}(P_{\mathrm{TF}} + P_{\mathrm{Target}} + N_{\mathrm{f}}) + k_{\mathrm{off}}(1 - P_{\mathrm{TF}}) \\
&\quad + k_{\mathrm{off,t}}(1 - P_{\mathrm{Target}}) + k_{\mathrm{off,d}}(N - N_{\mathrm{f}}), \\
\frac{dn_{\mathrm{Target}}}{dt} &= bm_{\mathrm{Target}} - \gamma n_{\mathrm{Target}}, \\
\frac{dP_{\mathrm{TF}}}{dt} &= -k_{\mathrm{on}} n_{\mathrm{TF}} P_{\mathrm{TF}} + (k_{\mathrm{off}} + \gamma)(1 - P_{\mathrm{TF}}), \\
\frac{dP_{\mathrm{Target}}}{dt} &= -k_{\mathrm{on}} n_{\mathrm{TF}} P_{\mathrm{Target}} + (k_{\mathrm{off,t}} + \gamma)(1 - P_{\mathrm{Target}}), \\
\frac{dN_{\mathrm{f}}}{dt} &= -k_{\mathrm{on}} n_{\mathrm{TF}} N_f + (k_{\mathrm{off,d}} + \gamma)(N - N_{\mathrm{f}}), \\
\frac{dm_{\mathrm{TF}}}{dt} &= r_0 P_{\mathrm{TF}} + r(1 - P_{\mathrm{TF}}) - \gamma_m m_{\mathrm{TF}}, \\
\frac{dm_{\mathrm{Target}}}{dt} &= r_{t,0} P_{\mathrm{Target}} + r_t(1 - P_{\mathrm{Target}}) - \gamma_m m_{\mathrm{Target}}.
\end{aligned} \tag{1}$$

Here, $P_{\mathrm{TF}}$, $P_{\mathrm{Target}}$ and $N_{\mathrm{f}}$ represent the free promoters of TF, target and free decoys, respectively and $m_{\mathrm{TF/Target}}, n_{\mathrm{TF/Target}}$ are the mRNA and protein for TF and target gene. The steady state expressions for TF and target gene can be obtained by setting the right hand side of the ODEs to zero and solving for each of the variables. Furthermore, using ODE solvers (ode45 in MATLAB or any other program) we can obtain both the steady state and the time to reach a certain threshold, we call it response time which is fundamentally different from the MFPT. Like mean first passage time (MFPT) from stochastic models, the response time from ODEs yield non-monotonic behavior for both auto-activating and auto-repressing genes (see Fig. 1 A-D). Also, the power-law behaviors obtained from deterministic model capture the same from stochastic simulations closely (Fig. 1 E,F)). However, the quantitative values between the two models differ depending on the parameter values. We have previously shown that depending on the network architecture the stochastic model and deterministic model can yield different quantitative outcomes [1].

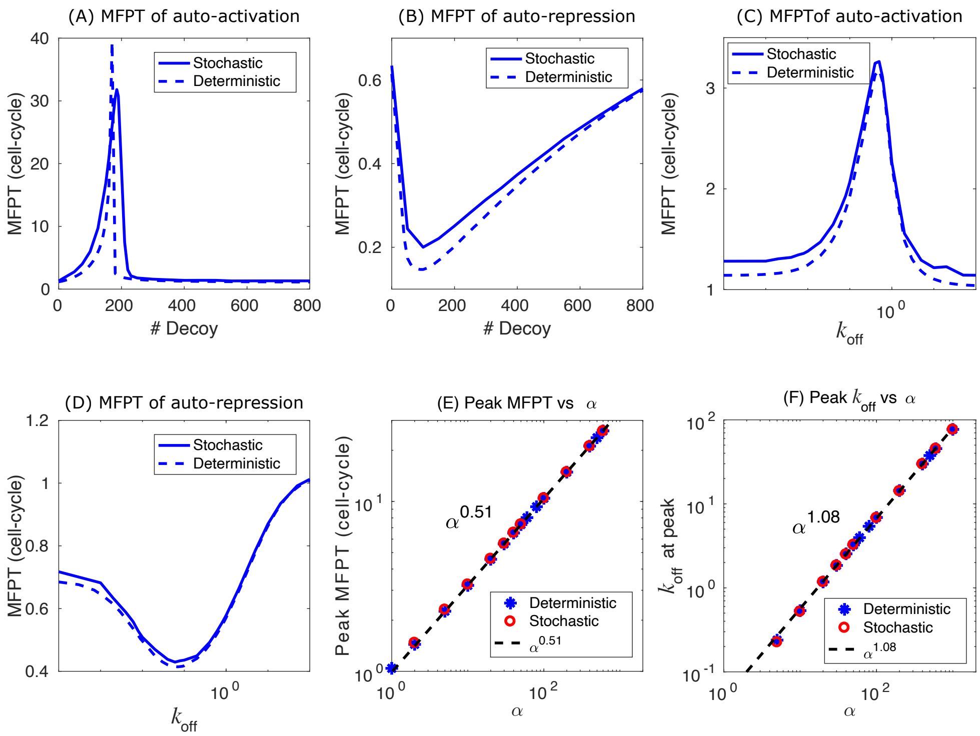


**Fig 1. Comparison of MFPT from stochastic simulation and deterministic simulation.** MFPT of an auto-activated (A) and and auto-repressed (B) TF gene when the number of decoy sites are tuned. (C,D) show the MFPT as a function of TF off-rate ($k_{\text{off}}$) for auto-activating (C) and auto-repressing TF gene (D). Solid and dashed lines correspond to the MFPT obtained using stochastic simulation and response time from deterministic model, respectively. (E) Parameters: $\alpha = 10$, $r_0 = 0.0025\ \text{s}^{-1}$, $b = 0.025\ \text{s}^{-1}$ (auto-activation), $\alpha = 0.1$, $r_0 = 0.005\ \text{s}^{-1}$, $b = 0.05\ \text{s}^{-1}$ (auto-repression). $\gamma_{\text{m}} = 0.01\ \text{s}^{-1}\text{mRNA}^{-1}$, and $\tau = 50$ min is used to generate all the panels. (E). Peak value of MFPT (or response time) as a function of auto-regulation strength, $\alpha$, when $k_{\text{off}}$ is tuned in deterministic model (blue asterisks) shows a power-law behavior with exponent similar to that of the stochastic model (red circles). (F) The values of $k_{\text{off}}$ at the peak MFPT as a function of $\alpha$ also capture the trend from stochastic model.

### References


1. Ali MZ, Parisutham V, Choubey S, Brewster RC. Inherent regulatory asymmetry emanating from network architecture in a prevalent autoregulatory motif. Elife. 2020;9.
