## Supplementary material for "Controlling gene expression timing through gene regulatory architecture": S3 Text

### S3 Text: Toy-model for power-law behavior

In order to determine the dependence of peak MFPT (response time for deterministic system) on regulatory strength ($\alpha$) and TF-binding rate ($k_{\mathrm{off}}$), we probe a simple toy model of auto-regulation. In this model, we ignore the mRNA dynamics and also consider the binding and unbinding events are fast enough to attain quasi-static equilibrium. If there are $x$ TF present at any time the probability that the gene is free ($P_{\mathrm{f}}$) or bound ($P_{\mathrm{b}}$) can be expressed as

$$P_{\mathrm{f}} = \frac{k_{\mathrm{off}}}{k_{\mathrm{on}}x + k_{\mathrm{off}}} = \frac{1}{\sigma x + 1}, \quad P_{\mathrm{b}} = \frac{k_{\mathrm{on}}x}{k_{\mathrm{on}}x + k_{\mathrm{off}}} = \frac{\sigma x}{\sigma x + 1}. \tag{1}$$

Here, $\sigma = k_{\mathrm{on}}/k_{\mathrm{off}}$. Now the rate of change of $x$ can be expressed as

$$\begin{aligned} \frac{dx}{dt} &= r_0 P_{\mathrm{f}} + r P_{\mathrm{b}} - \gamma x, \\ &= r_0 \frac{1 + \alpha\sigma x}{1 + \sigma x} - \gamma x. \end{aligned} \tag{2}$$

$r_0$ and $r$ being the basal production rate and production rate when TF is bound. The regulatory strength $\alpha = r/r_0$ is same as previously defined. The above differential equation can be solved exactly to obtain,

$$\begin{aligned} \ln(C) - \gamma\sigma t &= A\ln(x - a - b) + B\ln(x - a + b), \\ a &= \frac{r_0\alpha\sigma - \gamma}{2\gamma\sigma}, \quad b = \sqrt{a^2 + \frac{r_0}{\gamma\sigma}}, \\ A &= \frac{1 + \sigma(a + b)}{2b}, \quad B = \frac{-1 + \sigma(b - a)}{2b}. \end{aligned} \tag{3}$$

Here, $\ln(C) = A\ln(-a - b) + B\ln(-a + b)$ is a constant of integration obtained using initial condition, $x(t = 0) = 0$. Furthermore the steady state expression $x_{\mathrm{ss}}$ is given by $x_{\mathrm{ss}} = a + b$ for $t = \infty$. The response time ($\tau$), time to reach a certain fraction (f) of $x_{\mathrm{ss}}$, after doing a little bit algebra is expressed as

$$\tau = -\frac{\ln(1 - f)}{\gamma} + \frac{B}{\gamma\sigma}\left[\ln(1 - f) - \ln\big(1 + f\frac{a + b}{b - a}\big)\right]. \tag{4}$$

Eqn. 4, is a non-monotonic function with a complex dependence on the threshold fraction ($f$) and off-rate ($k_{\mathrm{off}} = k_{\mathrm{on}}/\sigma$). The first term in the expression is basically the response time for a constitutive gene, $\tau_{\mathrm{cons}} = -\ln(1-f)/\gamma = \ln(2)/\gamma$ for $f = 0.5$. In principle, to obtain peak response time, one can maximize eqn. 4. However, in this case, cannot be solved explicitly. In Fig. 1, we show the results obtained using Eqn. 4 which closely match with the full deterministic model.

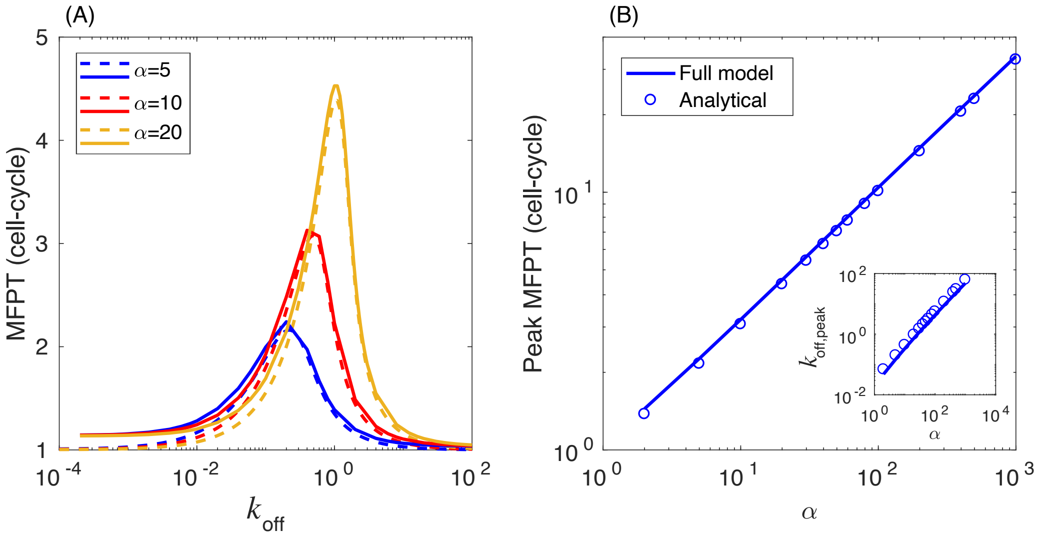


**Fig 1. Comparison of the full deterministic model and the simplified toy-model without mRNA dynamics.** (A) MFPT (response time) as a function of $k_{\mathrm{off}}$ from deterministic model (solid lines) and from Eqn. 4 for simplified toy-model for varying regulatory strengths ($\alpha$). (B) Peak MFPT versus $\alpha$ as well as $k_{\mathrm{off}}$ at peak versus $\alpha$ (inset) from the toy-model matches well with the full deterministic model.
